# Faster SARS-CoV-2 sequence validation and annotation for GenBank using VADR

**DOI:** 10.1101/2022.04.25.489427

**Authors:** Eric P Nawrocki

## Abstract

**Background:** In 2020 and 2021, more than 1.5 million SARS-CoV-2 sequences were submitted to GenBank. The initial version (v1.0) of the VADR (Viral Annotation DefineR) software package that GenBank uses to automatically validate and annotate incoming viral sequences is too slow and memory intensive to process many thousands of SARS-CoV-2 sequences in a reasonable amount of time. Additionally, long stretches of ambiguous N nucleotides, which are common in many SARS-CoV-2 sequences, prevent VADR from accurate validation and annotation.

**Results:** VADR has been updated to more accurately and rapidly annotate SARS-CoV-2 sequences. Stretches of consecutive Ns are now identified and temporarily replaced with expected nucleotides to facilitate processing, and the slowest steps have been overhauled using *blastn* and *glsearch*, increasing speed, reducing the memory requirement from 64Gb to 2Gb per thread, and allowing simple, coarse-grained parallelization on multiple processors per host.

**Conclusion:** VADR is now nearly 1000 times faster than it was in early 2020 for processing SARS-CoV-2 sequences submitted to GenBank. It has been used to screen and annotate more than 1.5 million SARS-CoV-2 sequences since June 2020, and it is now efficient enough to cope with the current rate of hundreds of thousands of submitted sequences per month. Version 1.4.1 is freely available (https://github.com/ncbi/vadr) for local installation and use.

## Background

The coronavirus disease 2019 (COVID-19) pandemic caused by the severe acute respiratory syndrome coronavirus 2 (SARS-CoV-2) has resulted in widespread illness, death and economic disruption [1, 2, 3]. The public health response to the pandemic has included efforts to sequence SARS-CoV-2 genomes from samples taken from infected individuals. The resulting nucleotide sequence data is deposited into databases where it can be accessed and analyzed, enabling researchers worldwide to study the variation present in the virus, understand its evolution, and monitor the spread of different variants in communities and countries around the world [4, 5, 6, 7, 8]. Validation of the sequence data prior to acceptance into the databases aides these analyses by limiting the amount of incorrect, artifactual sequence data they are based on. Annotation of biologically relevant features of the sequence data, such as protein-coding sequences and structural and functional RNA regions, provides additional relevant information.

The GenBank database maintained by the National Library of Medicine’s (NLM) National Center for Biotechnology Information (NCBI), is a comprehensive genetic sequence database in the public domain [9]. GenBank, along with the European Nucleotide Archive (ENA) [10] and the DNA Databank of Japan (DDBJ) [11] make up the International Nucleotide Sequence Database Collaboration (INSDC) [12] which was designed to allow free, unlimited and worldwide access to the most current and comprehensive set of DNA sequence information. In 2020 and 2021, over three million SARS-CoV-2 sequences were deposited into INSDC databases, including more than 1.5 million into GenBank. NLM’s NCBI also maintains the public Sequence Read Archive (SRA) database [13]. Data submitted to SRA typically includes sequence data from individual reads from sequencing runs, and data submitted to GenBank typically includes assembled consensus sequences that are full length or nearly full length SARS-CoV-2 genomes.

Due to the importance of SARS-CoV-2 data, GenBank created a dedicated and specialized submission portal workflow (https://submit.ncbi.nlm.nih.gov/sarscov2/) for data submitters to facilitate sequence and associated metadata upload [14]. Sequences submitted through this workflow are processed by an automated pipeline that performs quality control checks for potential sequencing artifacts and computes feature annotation, including protein coding sequences (CDS), mature peptide (mat peptide) and structural RNA features (stem loop) for each sequence. Sequences that meet all of the acceptance criteria are automatically deposited into GenBank, typically within minutes of submission. This automatic procedure for sequence validation and publication allows researchers to rapidly make their data publicly available for free, unrestricted use by the wider research community.

The GenBank SARS-CoV-2 submission portal pipeline first became available in June 2020, and since then the volume of SARS-CoV-2 sequence data submitted to GenBank has steadily increased. As shown in Table 1, by the end of 2020 about 43,000 SARS-CoV-2 sequences had been deposited into GenBank. In 2021, more than 1.5 million sequences were deposited, including more than 280,000 in December alone. This explosive growth in sequence data is primarily due to increased surveillance efforts by research groups around the world, including public health labs [15].

**Table 1.** Number of SARS-CoV-2 sequences released in GenBank in 2020 and 2021. Sequence counts were obtained using the NCBI Virus SARS-CoV-2 Data Hub on January 19, 2022, filtering by release date.

| month | year | #new<br>seqs | #cumulative<br>seqs |
| --- | --- | --- | --- |
| Jan | 2020 | 32 | 32 |
| Feb | 2020 | 58 | 90 |
| Mar | 2020 | 332 | 422 |
| Apr | 2020 | 1541 | 1963 |
| May | 2020 | 2974 | 4937 |
| Jun | 2020 | 3394 | 8331 |
| Jul | 2020 | 3604 | 11,935 |
| Aug | 2020 | 3818 | 15,753 |
| Sep | 2020 | 6731 | 22,484 |
| Oct | 2020 | 11,939 | 34,423 |
| Nov | 2020 | 4274 | 38,697 |
| Dec | 2020 | 4530 | 43,227 |
| Jan | 2021 | 8775 | 52,002 |
| Feb | 2021 | 26,078 | 78,080 |
| Mar | 2021 | 42,607 | 120,687 |
| Apr | 2021 | 97,095 | 217,782 |
| May | 2021 | 104,729 | 322,511 |
| Jun | 2021 | 46,187 | 368,698 |
| Jul | 2021 | 43,336 | 412,034 |
| Aug | 2021 | 141,958 | 553,992 |
| Sep | 2021 | 267,562 | 821,554 |
| Oct | 2021 | 239,296 | 1,060,850 |
| Nov | 2021 | 267,270 | 1,328,120 |
| Dec | 2021 | 288,771 | 1,616,891 |

### Viral sequence validation and annotation using VADR

The quality control sequence validation and annotation steps of GenBank’s automated pipeline are carried out by the VADR (Viral Annotation DefineR) software package [16], which performs multiple checks on each sequence and identifies the sequence coordinates of important features of the virus genome, specifically those features annotated in the SARS-CoV-2 NC_045512 RefSeq reference genome. This processing is performed by VADR’s *v-annotate*.*pl* program, which proceeds over four stages: *classification, coverage determination, alignment* and *protein validation*. Briefly, in the classification stage, for each sequence, the most similar reference model is determined and that model is used in the coverage determination stage to identify what regions of the sequence are similar to the reference and which are not. Typically the complete sequence will be similar to the reference, but potentially not if there is a misassembly or other problem. In the alignment stage, the entire sequence is then aligned to the best matching reference model and features are mapped from the model to the sequence based on that alignment. Finally, the coding potential of any predicted CDS regions are evaluated in the protein validation stage. For SARS-CoV-2 analysis since December 2, 2021, only one reference model, the NC 045512 RefSeq, is currently used, but multiple SARS-CoV-2 reference models were used previously, and multiple models are used for VADR processing of other viral sequences, including norovirus sequences.

At each stage, different types of unexpected situations can be detected and reported as *alerts* by the program. For example, one type of alert, with internal alert code *cdsstopn*, is reported if an early in-frame stop codon is detected in a predicted CDS region during the alignment stage, and another (*fstukcfi*) is reported if a potential frameshift is detected in a coding region. A subset of alerts, including *cdsstopn* and *fstukcfi*, are *fatal* in that they cause the sequence to *fail* and prevent it from automatic entry into GenBank. There are more than 70 different possible types of alerts, about 50 of which are fatal (see [16] and documentation in the package or on GitHub, e.g. https://github.com/ncbi/vadr/blob/master/documentation/alerts.md). VADR processing identifies potentially problematic sequences and information on failing sequences is sent back to submitters who are encouraged to check their data and resubmit with supporting information [14]. Output from VADR with detailed information on alerts, including relevant sequence and reference model coordinates, is provided to help in reviewing the sequences, but the high volume of SARS-CoV-2 data undoubtedly makes this process challenging for many submitters.

VADR was initially designed and tested on norovirus and dengue virus sequences which differ from SARS-CoV-2 sequences in several important ways that are relevant to VADR processing, as shown in Table 2. First, full length norovirus and dengue virus genome sequences are much shorter than SARS-CoV-2 sequences (about 7,500nt and 10,700nt, respectively versus about 29,900nt). Second, norovirus and dengue virus sequences exhibit significantly more sequence diversity than do SARS-CoV-2 sequences (at least at the time of writing). On average, norovirus genome sequences are about 82% identical within genogroups and dengue virus genome sequences are about 94% identical within serotypes whereas SARS-CoV-2 genomes are more than 99% identical. Third, SARS-CoV-2 sequences include a larger fraction of ambiguous N nucleotides (about 1.4% of all nucleotides) than do norovirus (0.5%) and dengue virus (0.2%) sequences, and more of those ambiguous nucleotides are present in contiguous stretches: more than 38% of SARS-CoV-2 sequences include at least one stretch of 50 or more consecutive Ns, but only 1.0% and 0.4% of norovirus and dengue virus sequences do. Fourth, there are many more SARS-CoV-2 sequences submitted to GenBank (more than 1.6 million in 2020 and 2021) than norovirus and dengue virus sequences (less than 200,000 total sequences in the database, less than 20,000 of which were deposited in 2020 and 2021).

**Table 2.** Attributes of norovirus, dengue virus, and SARS-CoV-2 sequences in GenBank and VADR 1.0 performance metrics. Only sequences deposited in GenBank were considered (ENA and DDBJ sequences were not included). ‘length’: average length of all RefSeq sequences for each virus. ‘# seqs’: total number of GenBank sequences for each virus, SARS-CoV-2 sequences limited to those with publication date in 2020 and 2021. Norovirus and dengue virus sequences were not limited by date; ‘% seqs full length’: percentage of sequences that are full length, defined as *>*= 95% the average length of shortest RefSeq sequence (minimum length 6952nt for norovirus, 10,117nt for dengue virus and 28,408nt for SARS-CoV-2); ‘% Ns’: percentage of total nucleotides in all sequences that are Ns; ‘% seqs with stretch of *>*= 50 Ns’: percentage of all sequences that have at least one stretch of 50 or more consecutive Ns; The remaining four rows pertain to single-threaded VADR 1.0 processing of all full length sequences: ‘average % identity’: the average of the average pairwise sequence identity in the multiple sequence alignments, one per RefSeq-based model, created by VADR; ‘seconds per sequence (VADR 1.0)’: the average running time per sequence (seconds); ‘required RAM (VADR 1.0)’: amount of RAM required; ‘total running time, CPU days (VADR 1.0)’: the total number of CPU days required; CPU times were measured as single threads on 2.2 GHz Intel Xeon processors. List of all norovirus and dengue virus sequences obtained by the following Entrez nucleotide queries on January 25, 2022, and then restricting to only GenBank sequences: “Norovirus NOT chimeric AND 50:10000[slen]” and “Dengue NOT chimeric AND 50:11200[slen]”; List of all SARS-CoV-2 sequences obtained using the NCBI Virus SARS-CoV-2 dashboard tabular view, restricting “release date” to 2020 and 2021. SARS-CoV-2 VADR 1.0 running time and average percent identity statistics are based on only 300 randomly selected SARS-CoV-2 sequences to limit total running time. Additional details are available in the supplementary material (https://github.com/nawrockie/vadr-sarscov2-paper-supplementary-material).

|  | *Norovirus* | *Dengue virus* | SARS-CoV-2 |
| --- | --- | --- | --- |
| length | 7575.6 | 10703.5 | 29903.0 |
| # seqs | 44,936 | 113,211 | 1,616,891 |
| % seqs full length | 5.1% | 8.4% | 99.7% |
| % Ns | 0.5% | 0.2% | 1.4% |
| % seqs with stretch of $\geq 50$ Ns | 1.0% | 0.4% | 38.7% |
| average % identity | 81.6% | 94.4% | 99.4% |
| seconds per sequence (VADR 1.0) | 42.4 | 92.6 | 331.8 |
| required RAM (VADR 1.0) | 8Gb | 8Gb | 64Gb |
| total running time, CPU days (VADR 1.0) | 1.1 | 10.2 | 6187.6 |

These differences have important implications on the practicality of processing SARS-CoV-2 sequences using the version of VADR (v1.0) available in early 2020 when the COVID-19 pandemic started. The relatively long length of the genome means that the classification, coverage determination and alignment stages are too slow for practical use on thousands of sequences because the running time of those stages empirically scale with the square of the genome length or worse. Processing a single full length SARS-CoV-2 sequence using VADR 1.0 takes approximately five minutes and processing all GenBank SARS-CoV-2 sequences from 2020 and 2021 would take more than 6100 CPU days (about 17 CPU years, Table 2). Additionally, the amount of memory required for VADR to process 30kb sequences is very high at 64Gb, meaning that parallelization on multiple processors is impractical because each processor would require 64Gb of RAM. Further, VADR v1.0 has difficulty accurately validating and annotating sequences with long stretches of Ns, which many SARS-CoV-2 sequences have, and such sequences often fail because of alerts related to low sequence similarity with the reference model due to the Ns.

### Implementation

I addressed the issues with SARS-CoV-2 sequence processing by implementing the following changes in VADR’s *v-annotate*.*pl* program:

- To accelerate processing of SARS-CoV-2 sequences, the slowest steps were modified to use *blastn* [17] (enabled with the `-s` option).
- To reduce the amount of memory required and to enable further acceleration via multi-threading, the memory expensive *cmalign* program was replaced with the memory efficient and faster *glsearch* program from the FASTA package [18, 19] for the alignment stage (enabled with the `--glsearch`, `--split` and `--cpu` options).
- To better handle long stretches of Ns, a pre-processing step was added that identifies regions of Ns and, when appropriate, temporarily replaces them with the expected nucleotides from the reference model sequence for purposes of validation and annotation (enabled with the `-r` option).

More detail on each of these changes is provided below.

### Improving speed and memory efficiency using blastn and glsearch

The classification and coverage determination stages of VADR annotation are typically performed by two separate rounds of the *cmsearch* program from the Infernal software package [20, 21], first in a faster score-only mode for classification and again in a slower mode that returns scores and model boundaries of hits to be used for coverage determination. When the *v-annotate*.*pl* `-s` option is used, *blastn* is executed once to determine the best matching model (subject) as well as the boundaries of all hits to that model. This single *blastn* step is roughly 200 times faster than the two *cmsearch* runs for typical SARS-CoV-2 sequences. The more similar the input sequence is to the reference model sequence, the less likely it is that replacing *cmsearch* with *blastn* will change the resulting alerts or annotation, and so this approach is well-suited for typical SARS-CoV-2 sequences which are highly similar to the NC 045512 reference.

The sequence alignment in the alignment stage of VADR v1.0 is performed by Infernal’s *cmalign* program. For SARS-CoV-2 (size 30kb), the amount of RAM required for *cmalign*, when using its default banded dynamic programming algorithm [22], can be up to 64Gb, and alignment typically takes about four minutes per sequence. With the `-s` option, this stage is modified to use the top scoring *blastn* alignment already computed from the classification and coverage determination stage. An alignment *seed* derived from that *blastn* alignment becomes fixed and only sequence regions outside the seed region (if any) are aligned in a separate step. In general, the more similar each input sequence is to the reference model, the longer the seed alignment will be, making this approach well suited for typically highly similar SARS-CoV-2 sequences. The seeded alignment strategy is depicted in Figure 1.

**Figure 1.**
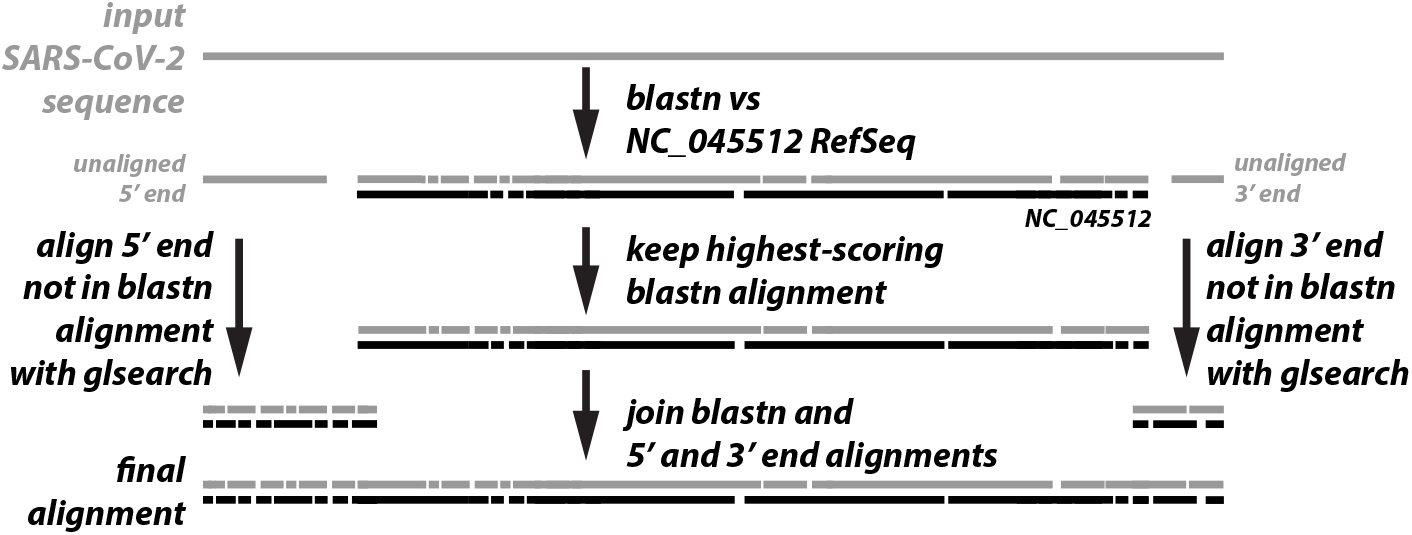
Seeded alignment strategy. The input sequence is used as a *blastn* query against the NC 045512 RefSeq sequence and the top-scoring alignment is kept as the seed, after potentially shortening it as described in the text. The 5’ and/or 3’ regions not covered by the seed (if any) plus 100 nucleotides of flanking sequence, are aligned to the full NC 045512 sequence using *glsearch*, and the resulting alignments are joined with the seed to produce the final alignment.

If the seed alignment includes the entire sequence, then the alignment stage is complete. This is the case for about 96% of sequences from 2020 and 2021 (Table 3).

**Table 3.** Summary statistics for N-replacement and seeded alignment on ENA SARS-CoV-2 sequences. The four rightmost columns report statistics for VADR 1.4.1 processing of subset of sequences used for testing (14,912 total, described in text). ‘seed coverage’ is defined as the percentage of the input sequence that is contained within the *blastn*-derived seed, if seed coverage is 100% this means that no subsequence alignment with *glsearch* was necessary. *v-annotate*.*pl* command line options used: `--cpu 8 --split --mdir <model-path> -s -r --nomisc --mkey sarscov2 --lowsim5seq 6 --lowsim3seq 6 --alt_fail lowscore,insertnn,deletinn --glsearch`, where `<model-path>` is the local path to the VADR SARS-CoV-2 v1.3-2 model set, matching current GenBank pipeline usage of VADR v1.4.1. Additional details are available in the supplementary material (https://github.com/nawrockie/vadr-sarscov2-paper-supplementary-material).

| month | year | # seqs<br>total | # seqs<br>tested | average<br># Ns<br>per seq | average<br>% Ns<br>replaced | average<br>seed %<br>coverage | % seqs w/<br>100% seed<br>coverage |
| --- | --- | --- | --- | --- | --- | --- | --- |
| Jul | 2020 | 1067 | 1000 | 813.9 | 99.3% | 100.0% | 96.7% |
| Aug | 2020 | 667 | 667 | 290.1 | 96.9% | 99.0% | 96.9% |
| Sep | 2020 | 790 | 790 | 189.3 | 99.5% | 100.0% | 98.6% |
| Oct | 2020 | 517 | 517 | 26.5 | 98.5% | 100.0% | 97.5% |
| Nov | 2020 | 545 | 545 | 1108.4 | 98.6% | 99.9% | 96.1% |
| Dec | 2020 | 140 | 140 | 52.5 | 99.7% | 100.0% | 95.7% |
| Jan | 2021 | 1688 | 1000 | 4220.9 | 99.8% | 97.6% | 87.9% |
| Feb | 2021 | 253 | 253 | 3318.9 | 99.6% | 100.0% | 98.4% |
| Mar | 2021 | 2198 | 1000 | 1250.3 | 99.4% | 99.9% | 98.5% |
| Apr | 2021 | 116,569 | 1000 | 138.4 | 99.5% | 99.8% | 98.1% |
| May | 2021 | 112,618 | 1000 | 196.8 | 98.5% | 99.6% | 94.3% |
| Jun | 2021 | 200,490 | 1000 | 193.2 | 96.9% | 99.5% | 94.6% |
| Jul | 2021 | 171,287 | 1000 | 307.2 | 98.8% | 99.9% | 97.7% |
| Aug | 2021 | 116,116 | 1000 | 307.7 | 98.7% | 99.7% | 97.2% |
| Sep | 2021 | 165,371 | 1000 | 279.8 | 99.1% | 99.8% | 97.0% |
| Oct | 2021 | 157,664 | 1000 | 480.7 | 99.0% | 99.8% | 97.3% |
| Nov | 2021 | 170,434 | 1000 | 347.5 | 98.1% | 99.8% | 97.2% |
| Dec | 2021 | 188,246 | 1000 | 259.9 | 97.9% | 99.8% | 96.9% |
| all combined |  | 1,406,660 | 14,912 | 711.1 | 99.3% | 99.6% | 96.4% |

If not, any sequence outside the seed region is then realigned to the model using the *glsearch* program of the FASTA package [18, 19] which aligns globally with respect to the input sequence and locally with respect to the reference model. Up to two alignments are required. If the seed does not begin at sequence position 1, a *glsearch* alignment for the 5’ end of the sequence with 100nt overlap with the 5’ end of the seed is computed. Analogously, if the seed does not end with the final sequence position, a *glsearch* alignment for the 3’ end of the sequence with 100nt overlap with the 3’ end of the seed is computed.

If the *glsearch* and seed alignments are determined to be consistent, the fixed seed alignment is then joined with the *glsearch* alignments of the 5’ and/or 3’ ends by simply concatenating the alignments after removing the overlapping regions from the seed alignment. The 5’ subsequence *glsearch* alignment and seed alignment are considered consistent if the 3’-most overlapping nucleotide of the 5’ subsequence aligns to the same model position in both the *glsearch* and seed alignment. And, likewise, the 3’ subsequence *glsearch* alignment and seed alignment are considered consistent if the 5’-most overlapping nucleotide of the 3’ subsequence aligns to the same model position in both the *glsearch* and seed alignment. If either 5’ or 3’ *glsearch* alignments are not consistent with the seed alignment, a non-fatal *unjoinbl* alert is reported for the sequence and then the entire sequence is realigned using *glsearch*. In practice, this is rare, occurring in 6.6% of sequences for which *glsearch* is used, and less than 0.3% of total sequences in the ENA test set of 14,912 sequences from Table 4.

**Table 4.** VADR running time on SARS-CoV-2 sequences. Running times computed for the set of 14,912 ENA sequences tested in Table 3. Columns 2, 3, and 4 indicate whether ‘seeded alignment’ (`-s` option in versions 1.1 and later), ‘N replacement’ (`-r` option in versions 1.1 and later), and ‘glsearch’ (`--glsearch` option in versions 1.2 and later), were enabled (‘+’) or not used (‘ *−* ‘). ‘# cpus’ indicates number of CPUs that were run in parallel (`--cpu <n>` option in version 1.2 and later). For v1.4.1 rows, `--split` was also enabled. The v1.4.1 row with 8 CPUs is in bold because that configuration is currently used for GenBank processing. For v1.0 and v1.1 timings, the 14,912 sequences were divided into smaller input files of 150 sequences each, run separately, and timings were combined. For v1.4.1 timings, a single file of all sequences was used as input. v1.0 *v-annotate*.*pl* command line options used: `--mxsize 64000 -m <model-path>/NC_045512.cm -b <model-path> -i <model-path>/NC_045512.minfo`, where `<model-path>` is the local path to the VADR coronaviridae 55-1.0.2dev-5 model set; v1.1 *v-annotate*.*pl* command line options used: `-s –r --mxsize 64000 -m <model-path>/NC_045512.cm -x <model-path> -I <model-path>/NC_045512.minfo -n <model-path>/NC_045512.fa`, where `<model-path>` is the local path to the VADR coronaviridae v1.1-1 model set; v1.4.1 *v-annotate*.*pl* command line options used: `--cpu <n> --split --mdir <model-path> -s -r --nomisc --mkey sarscov2 --lowsim5seq 6 --lowsim3seq 6 --alt_fail lowscore,insertnn,deletinn --glsearch`, where `<n>` is replaced by value from “# cpus” column and `<model-path>` is the local path to the VADR SARS-CoV-2 v1.3-2 model set, matching current GenBank pipeline usage of VADR v1.4.1; Additional details are available in the supplementary material (https://github.com/nawrockie/vadr-sarscov2-paper-supplementary-material).

| VADR version | seeded align-ment? | N replace-ment? | glsearch? | # cpus | required RAM | secs per seq | hours per 100K seqs | speedup vs v1.0 |
| --- | --- | --- | --- | --- | --- | --- | --- | --- |
| v1.0 | — | — | — | 1 | 64 Gb | 329.91 | 9164.3 | — |
| v1.1 | + | + | — | 1 | 64 Gb | 49.35 | 1370.7 | 6.7 |
| v1.4.1 | + | + | + | 1 | 2 Gb | 2.51 | 69.8 | 131.4 |
| v1.4.1 | + | + | + | 2 | 4 Gb | 1.49 | 41.5 | 220.8 |
| v1.4.1 | + | + | + | 4 | 8 Gb | 0.65 | 18.0 | 509.9 |
| **v1.4.1** | + | + | + | **8** | **16 Gb** | **0.33** | **9.3** | **986.8** |
| v1.4.1 | + | + | + | 16 | 32 Gb | 0.23 | 6.5 | 1417.9 |
| v1.4.1 | + | + | + | 32 | 64 Gb | 0.13 | 3.7 | 2462.2 |

The alignment seed is defined as the top-scoring *blastn* alignment, which can include gaps (either an insertion or deletion in the sequence relative to the reference model), where each ungapped region of sequence between two gaps is referred to as an ungapped segment below. This seed is shortened in some situations. If the seed includes a gap at a position that is within a CDS start or stop codon in the reference, then the seed is shortened to its longest ungapped segment. This shortening is performed because *blastn* tends to place gaps differently than *glsearch* does in some situations that lead to unnecessary downstream fatal alerts if those gaps are in start or stop codons. If no gaps in the seed occur within start or stop codons, the seed is shortened by removing 5’- and 3’-terminal ungapped segments from both ends until the following two criteria are met:

1. the 5’- and 3’-most terminal ungapped segments either include the sequence terminus (the first position on the 5’ end, or the final position on the 3’ end) or have a length of at least 100nt, which corresponds to the overlap length between the seed and *glsearch* regions.
2. *all* ungapped segments have a minimum length of 10nt.

Criteria 1 is enforced because if a terminal segment has a length less than 100nt then there will almost certainly be a gap in the overlapping region of the seed and terminal aligned regions, which will increase the chance of an *unjoinbl* alert triggering the relatively slow step of *glsearch* realignment of the full sequence. Criteria 2 is enforced because in gap-rich regions *glsearch* creates alignments that are less likely than *blastn* alignments to result in unnecessary downstream alerts (based on internal testing, not shown).

### Handling N-rich regions

As shown in Table 2, N-rich regions are relatively common in SARS-CoV-2 sequences. An ambiguous N character is used in nucleotide sequence data to indicate that the identity of the corresponding nucleotide in the sequence is unknown, and Ns are permitted in GenBank SARS-CoV-2 sequence data if they are the appropriate length with respect to the NC_045512 reference sequence. A region of Ns should not, for example, introduce a frameshift that changes the reading frame of a proteincoding region (CDS feature). But because VADR validates and annotates sequences based on similarity to a reference model, regions of ambiguous Ns often cause sequences to fail that should pass, preventing them from automatic publication in GenBank.

To address this, *v-annotate*.*pl* has an optional pre-processing step enabled with the `-r` option that identifies stretches of Ns and temporarily replaces them, if possible, with the expected nucleotides from the homologous positions in the reference model sequence. The validation and annotation of the sequence then proceeds with the modified sequence. If the sequence passes, the original sequence that includes Ns is deposited into GenBank, and feature annotation is “trimmed” to indicate which features begin and/or end with one or more Ns. Trimming involves changing the coordinates of the feature start/stop positions to the first/final non-ambiguous nucleotide. The N-replacement strategy is depicted in Figure 2.

**Figure 2.**
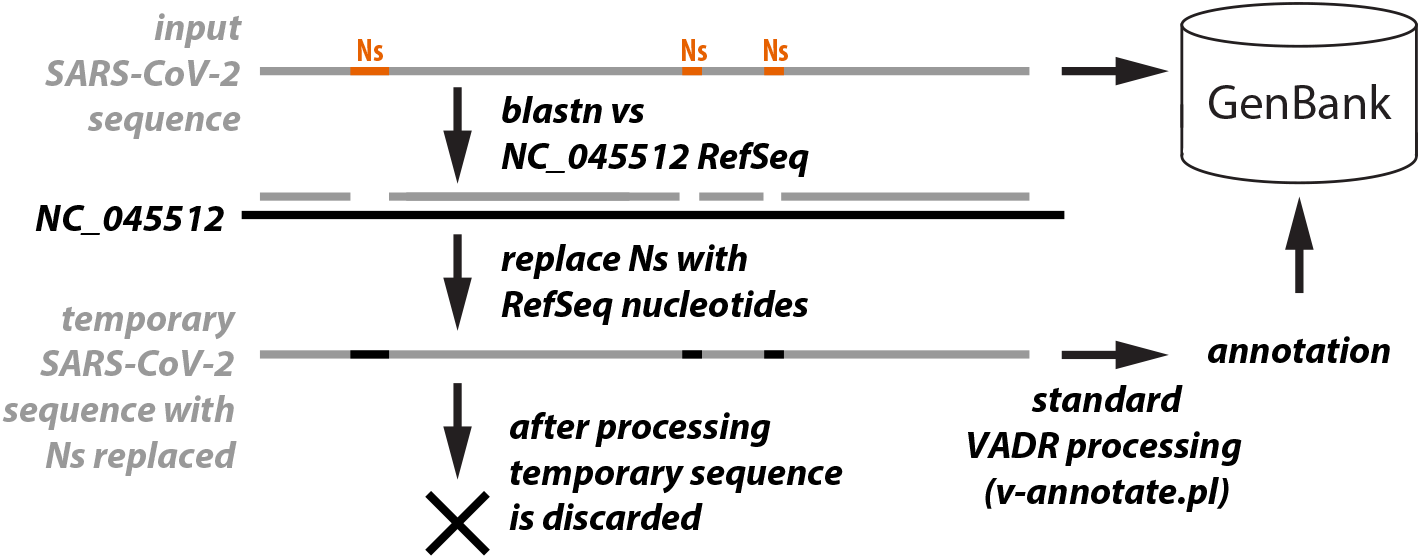
N-replacement strategy. Stretches of contiguous Ns in input sequences (orange) are identified as regions not covered by *blastn* alignments (gaps in second grey line) after the input sequence is used as a query against the NC_045512 RefSeq sequence. The Ns in these regions are replaced by the corresponding ‘expected’ nucleotides from NC_045512 and the resulting sequence is validated and annotated by *v-annotate*.*pl*. Importanty, the sequence deposited into GenBank is the original input sequence, including the Ns, not the sequence with the Ns replaced that was processed by *v-annotate*.*pl*.

The N-replacement step begins by running *blastn* for each input sequence. All hits are then sorted by input sequence start position. In the rare situation that one hit is completely spanned by another, the lower scoring hit is discarded. The missing regions in both the input sequence and reference model sequence not covered by any of the non-discarded hits are then evaluated to determine if they satisfy the criteria listed below. The evaluation of each missing region between two hits involves comparing the missing sequence region *S* with the corresponding missing reference model region *M*. *S* and *M* are checked to see if:

1. the length of *S* is at least 5 nucleotides.
2. the fraction of N nucleotides in *S* is at least 0.25 if *S* contains the first or final nucleotide in the sequence, or at least 0.5 if it does not.
3. *S* and *M* are equal length. Command line options exist to change the thresholds in criteria 1 (`--r_minlen`), and criteria 2 (`--r_minfract5`, `--r_minfract3`, and `--r_minfracti`). In December 2021, it became clear that some N-rich SARS-CoV-2 omicron variant sequences failed VADR v1.3 due to low similarity with the model because some N-rich regions that satisifed criteria 1 and 2 but not 3 were not being replaced. This tended to occur with N-rich regions close to omicron-specific insertions or deletions relative to the NC_045512 reference model. It is reasonable to assume that such insertions and deletions will become more common as SARS-CoV-2 continues to mutate and new variants arise. This situation motivated a change in VADR v1.4 to allow Ns to be replaced even if criteria 3 is not met if the following additional criteria are met:
4. difference in length between the *S* and *M* is at most 10nt
5. *S* includes at least 1 nucleotide that is *not* an N
6. the fraction of non-N nucleotides in *S* that are identical to the expected nucleotides in *M* is at least 0.75

Command line options exist to change the thresholds in criteria 4 (`--r_diffmaxdel`, and `--r_diffmaxins`), criteria 5 (`--r_diffminnonn`) and criteria 6 (`--r_diffminfract`).

Because *S* and *M* are different lengths, the two regions must be aligned in some way to determine which positions of *S* correspond to which positions of *M* in order to compare non-N nucleotides in *S* with the expected nucleotides in *M* when checking criteria 6. Two simple alignment strategies are attempted: (i) a 5’-flush alignment in which the first nucleotide of *S* is aligned to the first nucleotide of *M* and a gapless alignment is extended towards the 3’ end for *L* positions (where *L* is the shorter of the lengths of *S* and *M*) with any extra positions in either *S* or *M* aligned to gaps at the 3’ end of the alignment; and (ii) a 3’-flush alignment in which the final nucleotide of *S* is aligned to the final nucleotide of *M* and a gapless alignment is extended towards the 5’ end for *L* positions with any extra positions in either *S* or *M* aligned to gaps at the 5’ end of the alignment. The fraction of matches between observed non-N nucleotides in *S* and expected nucleotides in *M* is determined for the 5’-flush and 3’-flush alignments and if either is at least 0.75, then criteria 6 is met.

When criteria 1, 2 and 3 are met, any N nucleotides in *S* are replaced by the corresponding expected nucleotides in *M* (the correspondence is trivial to determine because *S* and *M* are the same length, effectively implying a gapless alignment).

Alternatively, when criteria 3 is not met but criteria 4, 5 and 6 are met, any N nucleotides in *S* are still replaced by the corresponding expected nucleotides in *M* but the correspondence is not trivial due to the difference in length between *S* and *M*. In this case, if the 5’-flush ‘alignment’ yielded a higher (or equal) fraction of matches in non-N nucleotides between *S* and *M*, then that alignment is used to determine the corresponding nucleotides for replacement of Ns. If the 3’-flush alignment yielded a higher fraction of matches, then that alignment is used.

### Coarse-grained parallelization for faster processing

VADR processes each sequence independently, so splitting up the sequence file into multiple partitions and processing each partition independently on a single CPU is a simple and effective parallelization strategy on multi-processor machines. With version 1.0, this strategy was impractical on typical hardware because the alignment stage would have required up to 64Gb of RAM per CPU and so would have only been possible on extremely high memory host machines. Using the memory-efficient *glsearch* program and making sufficiently small partitions in version 1.4.1, the memory requirement is reduced to 2Gb per CPU in practice making parallelism across multiple CPUs practical. When run with the `--split --cpu <n>` options, *v-annotate*.*pl* runs in a special mode that first splits the input sequence file into *x* smaller files, each with about 10 SARS-CoV-2 sequences (total length of 300kb), and processes each file independently with a separate execution of *v-annotate*.*pl* (this time without the `--split` and `--cpu` options) on `<n>` CPUs in parallel requiring a total of `<n>`*∗*2Gb of total RAM on the host machine. When all of the sequences have been processed, the output from the *x* runs of *v-annotate*.*pl* are then merged into the final output files. The final output is identical to what would be obtained if the program was run without parallelization on the full input sequence file (e.g. without the `--split` and `--cpu` options). Because information on each sequence (e.g. annotations, alerts) is kept in memory by the program, the amount of RAM required for this information scales with the total length of all sequences in the input sequence file. This is why the size of the *x* files is limited to 300kb, which empirically reduces the required memory to 2Gb RAM per CPU. The final step that merges the output of the *x* runs is performed in a memory efficient manner that does not require keeping data from all sequences in memory. In practice, GenBank uses the `--split --cpu 8` options when running *v-annotate*.*pl* to parallelize over 8 CPUs.

## Results

To characterize the efficacy of the N-replacement and seeded alignment strategies, VADR v1.4.1 was tested on a random sample of 14,912 SARS-CoV-2 sequences deposited in the the ENA database between July 1, 2020 and December 31, 2021.This dataset is composed of 1000 randomly selected sequences from each month in which at least 1000 were published in ENA, and all sequences from every other month. Only four SARS-CoV-2 sequences were published in ENA prior to July 2020 and these were not included. ENA sequences were chosen for testing because all GenBank sequences have been pre-screened by VADR, whereas ENA does not include VADR as part of its submission pipeline, and so ENA sequences represent a more realistic sample of submitted sequences that would be input to *v-annotate*.*pl* in the GenBank submission portal pipeline. Terminal ambiguous nucleotides including Ns at the beginning and end of the sequences were removed prior to testing, such that the first and final nucleotide of each sequence are non-ambiguous, because the GenBank submission portal trims sequences in this way prior to VADR processing. Table 3 summarizes characteristics of the N-replacement and seeded alignment strategy on this dataset. Each sequence contains on average about 711 Ns, 99.3% of which are replaced using the *blastn*-based N-replacement strategy. After N-replacement, the alignment seed covers 99.6% of each sequence on average, and is the complete sequence for 96.4% of sequences. The statistics are generally consistent over the 18 month period, with only slight month-to-month variability.

Table 4 summarizes the impact of the N-replacement, seeded alignment and parallelization strategies implemented in VADR v1.4.1 on the processing time for SARS-CoV-2 sequences compared with v1.0 and v1.1. Version 1.0 did not include any of the acceleration strategies discussed here. Version 1.1 included a simplified version of both N-replacement (only criteria 1, 2 and 3 above were enforced) and seeded alignment (the seed was defined as the longest single ungapped region of the top scoring *blastn* alignment) but used *cmalign* instead of *glsearch* to align regions outside the seed and did not allow multi-CPU parallelization.

The N-replacement and seeded alignment strategies alone offered about a 7-fold speedup in v1.1 versus v1.0, but did not lower the high 64Gb RAM requirement. In v1.4.1, combining those two strategies with *glsearch* and the `--split` option on a single CPU yielded about a 130-fold speedup relative to v1.0 and reduced the required RAM to 2Gb. Parallelization on multiple CPUs using the `--cpu` option resulted in further acceleration, decreasing the number of seconds per sequence to 0.13 seconds (2400-fold speedup vs v1.0) when run on 32 CPUs. The GenBank submission portal currently runs VADR v1.4.1 on 8 CPUs in parallel, which processes SARS-CoV-2 sequences nearly 1000 times faster than VADR v1.0 did in early 2020. These data show that the combination of N-replacement and seeded alignment is highly effective on SARS-CoV-2 sequences from 2020 and 2021.

The data in Table 4 demonstrate the impact of N-replacement and seeded alignment on running time but not on pass/fail outcomes or annotation. If v1.0, which doesn’t use N-replacement, seeded alignment or *glsearch*, is used to process the 14,912 sequences, only 3892 sequences pass and 11,020 fail. If v1.4.1 is used with only N-replacement (`-r` option), 13,436 sequences pass and 1,576 sequences fail, indicating the importance of N-replacement on allowing N-rich sequences to pass. Alternatively, if v1.4.1 is run as recommended, using N-replacement, seeded alignment, and *glsearch* (`-r`, `-s`, and `--glsearch` options), 13,430 sequences pass (a subset of the 13,436), and 1,582 sequences fail. Of the six additional sequences that fail, four fail due to low similarity in the first 10 nucleotides (LR899087.1, LR992006.1, HG994328.1, HG994349.1) one fails due to a short potential frameshift (OU218955.1), and the other fails due to uncertainty in a mature peptide boundary (OU923822.1). All six of these cases are arguably acceptable failures as they are situations that warrant manual examination prior to publication in GenBank. Of the other 14,906 sequences with the same pass/fail outcomes between v1.4.1 with recommended options and v1.4.1 with only N-replacement, only 42 (0.3%) differ in terms of annotation of one or more features, often by a single nucleotide at the start or stop boundary of a single feature. The low number of differences in pass/fail outcomes (6/14,912, 0.04%) and feature annotations show that the seeded alignment and *glsearch* acceleration strategies have only a small impact on pass/fail outcomes and annotation.

## Discussion

GenBank began using VADR in 2018 for automatic validation and annotation of norovirus sequences [16], and later began applying it to dengue virus and SARS-CoV-2 sequences. The improvements to the efficiency of VADR for SARS-CoV-2 processing described here allowed GenBank to keep up with the dramatic increase in sequence submission numbers observed in 2020 and 2021. One alternative to using VADR would be for GenBank to not use software to screen incoming SARS-CoV-2 sequences and instead attempt a more manual review process, or even accept them without validating or annotating them. Manual review of all sequences is impractical for SARS-CoV-2 given the sheer volume of data and limited personnel at GenBank. Accepting data blindly as submitted would mean that sequences with sequencing artifacts and other problems, which VADR would have identified and prevented the publication of, would exist in the database and introduce noise into downstream analyses. This lack of validation would also presumably lower user trust in the integrity of the data and would mean that VADR-based annotation information of different sequence features, which increases usability and facilitates some types of analysis, would not exist. On the other hand, using VADR potentially prevents some legitimate sequences without errors to be rejected and not published in GenBank, especially if the submitter does not follow-up after the sequences are not accepted.

Because the growth in sequence data from tens of sequences a month to thousands per month occurred relatively quickly over the course of just a few months in early 2020 (Table 1), the speed improvements in VADR needed to be implemented, tested and deployed relatively quickly as well. The specifics of the changes described here were partly motivated by the fact that they could be implemented fast with modifications to the existing VADR codebase, as opposed to as part of a newly written program designed specifically for SARS-CoV-2.

Though effective, the N-replacement strategy is complex and not perfect as it does not replace all Ns in sequences (missing about 0.7% in the test dataset in Table 3). One alternative would be to not replace Ns but to instead modify the alignment and protein validation stages to better tolerate Ns, possibly by allowing Ns to align with any canonical nucleotide with a positive score. Unfortunately, this would require reimplementation of the core algorithms used by VADR, presently implemented in the efficient programs *blastn, glsearch* and *blastx*. Reimplementations of the tasks performed by these programs would be burdensome, and unlikely to match the efficiency of the relatively optimized existing implementations, likely slowing down processing. However, as SARS-CoV-2 continues to mutate, the current N-replacement strategy may prove too ineffective for continued use and need to be modified or overhauled, potentially by a radical change such as better tolerance of Ns instead of replacement. One obvious way the current strategy could be modified is if a more accurate alignment of the missing sequence and model regions were computed when they differ in length, using a Needleman-Wunsch-like algorithm [23].

The seeded alignment strategy performs well for existing SARS-CoV-2 sequences, but may need to be updated or rethought in the future. The effectiveness of the technique depends on the length of the best *blastn* alignment, and as divergence in SARS-CoV-2 sequences continues, the expected alignment length will decrease, lowering the impact of this approach and increasing running time. Depending on the practical implications of this on GenBank processing, it may be necessary to modify the seeded alignment strategy and/or introduce additional reference SARS-CoV-2 models for specific variants to address sequence divergence. Removing the *blastn*-based seeded strategy and instead using *glsearch* to align the full input sequences is likely not a viable option as that step alone would require about 15 seconds per sequence, which is about six times longer than the current implementation takes to completely process a sequence.

Currently, N-replacement, seeded alignment, and *glsearch*-based alignment are only used in the SARS-CoV-2 GenBank processing pipeline and not employed for norovirus or dengue virus sequences. This is partly because processing speed is not as relevant for those viruses due to the smaller genome sizes and lower number of submitted sequences (Table 2). If sequence submissions of those viruses were to increase, or if VADR is adopted for automated processing of additional viral sequences with higher sequence submission volume, (e.g. Human Immunodeficiency Virus-1 (HIV-1)) we may explore extending these acceleration strategies beyond SARS-CoV-2 processing.

The acceleration heuristics described here were not the only changes made to VADR SARS-CoV-2 processing in 2020 and 2021. In total, 14 releases of VADR were made between January 2020 and January 2022 (v1.0.2 to v1.4.1). The VADR GitHub Wiki (https://github.com/ncbi/vadr/wiki/Coronavirus-annotation) has detailed instructions, including a tutorial, on using VADR for SARS-CoV-2 analysis, as well as information on when each version of VADR was used for automated GenBank sequence processing.

Between January and April 2020, three alternative reference model sequences were added to the set of VADR models, including one for each of the B.1.1.7 (alpha) and B.1.525 variants and one with a deletion at NC_045512 reference position 28254. These models were added to allow sequences with certain natural mutations relative to NC_045512 to pass that would have otherwise failed due to one or more fatal alerts if only the NC_045512 reference model was used. In February 2021, v1.1.3 of VADR introduced the capability of allowing certain fatal alerts for specific features to *not* cause a sequence to fail but rather to cause those features to be annotated as a miscellaneous feature (misc feature) instead. Since February 19, 2021 this strategy has been used for the ORF8 CDS feature. On August 5, 2021, it was extended to the ORF3a, ORF6, ORF7a, ORF7b and ORF10 CDS features, as well as the stem loop from NC_045512 positions 29728 to 29768. On December 2, 2021, GenBank processing reverted to using a single model because the common mutations that motivated the addition of the three alternative models would no longer cause a sequence to fail due to the miscellaneous feature failover, and because using the single NC_045512 model was generally leading to more accurate, consistent, and justifiable results on the breadth of sequence diversity that was being observed at the time. The results of all tests reported here were performed with the single NC_045512 model, but the heuristics work similarly for alternative models. In the future, as SARS-CoV-2 continues to undergo mutations, it may be necessary to complement the NC_045512 model with additional models, for example for novel variants. If that happens, the heuristics here should work well in principle but may need to be refined based on attributes of the new sequence data.

## Conclusion

The improvements to the speed and memory efficiency of VADR described here for SARS-CoV-2 processing were necessary for GenBank to keep up with the high volume of sequence submissions in 2020 and 2021. Processing the more than 1.5 million GenBank sequences with the version that existed in January 2020 (v1.0) would have required more than 15 CPU years. With the current implementation in the GenBank SARS-CoV-2 processing pipeline, VADR v1.4.1 running in parallel on 8 CPUs, the same set of sequences would require roughly one week to process, and this could be further sped up by additional parallelization with more CPUs and/or additional hosts. VADR is freely available (https://github.com/ncbi/vadr) for users to download and run locally, enabling them to screen their own data prior to submission to GenBank or for other purposes.

## Availability and requirements

Project name: VADR

Project home page: https://github.com/ncbi/vadr

Operating system(s): Linux, Mac OS X

Programming language: Perl

Other requirements: Bio-Easel v0.15, BLAST+ v2.12.0, HMMER v3.3.2, Infernal v1.1.4, FASTA v36.3.8h, Sequip v0.09

License: public domain

Any restrictions to use by non-academics: none

## Abbreviations

COVID-19: coronavirus disease 2019
SARS-CoV-2: severe acute respiratory syndrome coronavirus 2
NLM: National Library of Medicine
NCBI: National Center for Biotechnology Information
ENA: European Nucleotide Archive
DDBJ: DNA Databank of Japan
INSDC: International Nucleotide Sequence Database Collaboration
VADR: Viral Annotation DefineR
BLAST: Basic Local Alignment Search Tool
nt: nucleotides
kb: kilobase (1000 nucleotides)
misc feature: miscellaneous feature

## Declarations

### Ethics approval and consent to participate

Not applicable.

### Consent for publication

Not applicable.

### Availability of data and materials

All data generated or analyzed during this study are included in this published article, its supplementary material, or NCBI’s GenBank database. Code is available on github (https://github.com/ncbi/vadr). The supplementary material includes instructions for reproducing the data reported in the tables in the article (https://github.com/nawrockie/vadr-sarscov2-paper-supplementary-material).

### Author’s contributions

EPN conceived of the software improvements, wrote the code and wrote the paper.

## Acknowledgements

Model changes and miscellaneous feature failover decisions were made in conjunction with Linda Yankie, Vince Calhoun and Ilene Mizrachi who also tested VADR changes along with Beverly Underwood, Susan Storz, and Eneida Hatcher. Thanks to Sergiy Gotvyanskyy and Denis Sinyakov for incorporating VADR into the GenBank submission portal system, to Jesse Becker, Ron Patterson and the NCBI systems team for expert management of the compute farm, and to Linda Yankie, Vince Calhoun, Eneida Hatcher, Beverly Underwood and Ilene Mizrachi for helpful comments on the manuscript. This research was supported by the Intramural Research Program of the NIH, National Library of Medicine.

## Additional Files

Supplementary material available at: https://github.com/nawrockie/vadr-sarscov2-paper-supplementary-material.

